# RNaseH-based ribodepletion of total planarian RNA improves detection of longer and non-polyadenylated transcripts

**DOI:** 10.1101/2024.07.20.604429

**Authors:** Pallob Barai, Shishir Biswas, Prince Verma, Elizabeth M. Duncan

## Abstract

The overwhelming majority of RNA species isolated from cells or tissues using organic extraction are ribosomal RNAs (rRNA), whereas a relatively small percentage are messenger RNAs (mRNA). For studies that seek to detect mRNA transcripts and measure changes in their expression, this lopsided ratio of desired transcripts to undesired transcripts creates a significant challenge to obtaining sensitive and reproducible results. One method for improving mRNA detection is to selectively amplify polyadenylated (polyA) mRNA molecules when generating RNA-seq libraries, a strategy that is generally very successful in many species. However, this strategy is less effective when starting with total RNA from some species e.g., the planarian species *Schmidtea mediterranea* (S.med), as it generates libraries that still contain significant and variable amounts of rRNA reads. Further, commercially available ribodepletion kits do not efficiently deplete rRNAs from these samples because their sequences are divergent from mammalian rRNAs. Here we report a customized, optimized, and economical ribodepletion strategy than allows the generation of comprehensive RNA-seq libraries with less than one percent rRNA contamination. We show that this method improves transcript detection, particularly for those without polyA tails (e.g., core histones) and those that are relatively long (e.g., microtubule motor proteins). Using this custom ribodepletion approach, we also detected many transcripts that are not represented in the most recent set of S.med gene annotations, including a subset that are likely expressed transposable elements (TEs). To facilitate future differential expression analyses of these newly identified loci, we created both an annotation file of the new loci we identified and a bioinformatic pipeline for generating additional annotations from future libraries. As significant recent research shows that TE activation is regulated and functionally important, the resources provided here will provide a starting point for investigating such mechanisms in planarians and other species with less conserved rRNA sequences.

## Introduction

RNA sequencing (RNA-seq) is a powerful method with many useful applications (Wilhelm and landry 2009; Kim *et al*. 2015; Tiong *et al*. 2024). First, it is commonly used to assess differential gene expression between control and treatment samples, including those induced by RNA interference (RNAi), enzyme inhibitors, or metabolite analogs. Second, RNA-seq is used to generate species and/or context-specific transcriptomes and to inform the creation of gene model annotations for a related genome assembly (Guttman *et al*. 2010; Bussotti *et al*. 2016; Singh *et al*. 2017; Lawson *et al*. 2020). Moreover, these two applications are interdependent: to compare the relative number of gene-specific RNA-seq reads between samples, those reads must first be assigned to a defined genomic (or transcriptomic) locus (Love *et al*. 2014). It is therefore critical to build an accurate and comprehensive set of gene models and/or transcripts, as reads will not be counted if they do not map to a defined sequence.

In eukaryotic cells, over 80% of the total RNA is ribosomal RNA (rRNA), non-coding RNA molecules that partner with specific proteins to form the small and large ribosomal subunits (Ide *et al*. 2010). This presents a challenge when aiming to accurately and efficiently quantitate changes in the remaining 10-20% of RNA species, including mRNAs. To address this issue, one of two types of RNA-sequencing strategies are usually employed: 1) polyA-enrichment, which targets RNAs that are polyadenylated during processing i.e., most mRNAs and some non-coding RNAs (ncRNAs), and 2) ribodepletion, in which the ribosomal RNAs (rRNAs) are depleted by affinity capture or enzymatic degradation, leaving all the “other” RNAs behind for sequencing. In the planarian field, the large majority of RNA-sequencing studies use the polyA-enrichment strategy (Swapna *et al*. 2018; Neiro *et al*. 2022; Ivankovic 2023). Although these datasets have produced many informative findings, this approach is not as efficient in its enrichment of polyadenylated planarian transcripts as it is in other species: whereas polyA-enrichment from mammalian total RNA often generates RNA-seq libraries with < 1% rRNAs (Cui *et al*. 2010), planarian polyA RNA-seq libraries often have significantly higher and variable amounts of rRNA ranging from 5-30% (Kim *et al*. 2019b). However, it is unclear why this occurs.

To combat this problem, a previous study used biotinylated rRNA probes to selectively remove rRNA species from samples of total RNA extracted from planarian stem cells (Kim *et al*. 2019b). Although this method is very effective, biotinylated probes and magnetic beads are expensive additions to the costs of creating and sequencing RNA libraries. Here we describe an optimized RNaseH-based ribodepletion strategy that is cost-effective and can be customized to any species. We used this strategy to generate several comprehensive paired-end RNA-seq libraries from both uninjured and regenerating planarian tissues. In examining these datasets and comparing them to highly similar polyA-enriched RNA-seq libraries, we found that our RNaseH-ribodepletion libraries provide markedly better detection of specific types of transcripts, including those encoding core histones, longer transcripts, and a distinct set of uncharacterized RNAs. To define and characterize the latter, we created a bioinformatic pipeline that identifies loci with significant representation of mapped RNA-seq reads relative to background, including many that are not currently annotated on the most recent planarian (S.med) genome (Ivankovic 2023). We then validated and analyzed these new loci further and found that a major fraction of them are likely actively transcribed transposable elements (TEs), including both DNA and RNA subclasses. Overall, these data provide a method, a deep transcriptomic resource, and a useful bioinformatic tool that can be applied across different datasets, sample types, and species. It also uncovers several avenues for futures studies, including the potential characterization of new histone variants, noncoding RNAs, and actively expressed TEs.

## Materials and Methods

### RNA extraction and ribodepletion with RNaseH

Total RNA was extracted from whole planarian worms or regenerating tissue using TRIzol reagent according to the manufacturer protocol. After extracting RNA, it was further cleaned, concentrated, and size-selected for RNAs > 200nt using the Zymo Research RNA Clean & Concentrator-5 kit (Catalogue No. R1013). An RNase H mediated rRNA depletion protocol was adapted from methods developed for bacteria (Culviner *et al*. 2020) and human cells (Baldwin *et al*. 2021). Briefly 500 ng of size-selected total RNA was mixed with 1000ng of a species-specific rRNA probe pool (**Supplemental Table S1**) and hybridization buffer (final 50mM Tris HCl pH 7.5, 100mM NaCl, 20mM EDTA) in a final volume of 15µL. Probe-RNA mixtures were heated in a heat block at 95°C for 2 mins and then slowly cooled (-0.1°C/sec) to 65°C. The mixtures were then kept at 65°C additional 5 minutes while the RNaseH mixture was prepared: for each RNA sample, 3μL Lucigen Hybridase™ Thermostable RNase H enzyme (Catalogue No. H39500), 0.5μL 1M Tris-HCl pH 7.5, 0.2μL 5M NaCl, 0.4μL 1M MgCl2, and 0.9μL nuclease-free water (total 5μL) was mixed and preheated to 65°C. The 5μl preheated Hybridase RNase H reaction was then added to the sample mix and incubated at 65°C for 2.5 minutes. In-column DNase I treatment was then performed to degrade excess probes and genomic DNA; 30µL of prepared DNase I reaction mix (3μL TURBO™ DNase [ThermoFisher AM2239] + 5μL10X TURBO™ DNase Buffer + 22μL nuclease-free water) was added to the samples immediately and incubated at 37°C for 30min. The ribodepleted RNA was then cleaned using the Zymo Research RNA clean and concentrator-5 kit (Catalogue No. R1013) and RNA-seq libraries were prepared using the KAPA RNA hyper kit (Catalogue No. 08098107702) along with KAPA unique dual index adapters (Catalogue No. 8861919702) according to the manufacturer protocol.

### RNA sequencing, alignment to the S3h1 genome, and identification of putative ncRNA loci

RNA-seq libraries were shipped to the DNA Services Lab in the Roy J. Carver Biotechnology Center at the University of Illinois Urbana Champaign, checked for quality, and sequenced using paired-end sequencing technology on an SP flowcell of the Illumina NovaSeq 6000. The resulting sequences were then aligned to the S3h1 genome (Ivankovic 2023) using HiSat2 (Kim *et al*. 2019a). Transcripts per Million mapped reads (TPMs) for each gene model were calculated using Stringtie (Pertea *et al*. 2015). Library complexity was analyzed using the RNA-seq QC R package (https://rdrr.io/cran/RNAseqQC/).To identify loci with significant RNA-seq read signal but no overlap with existing high-confidence h1SMc gene models (Ivankovic 2023), MACS2 (Zhang *et al*. 2008) was used to call “peaks” of significant RNA-seq reads (from a batched alignment of all RH-ribodepleted paired-end libraries; see Bioproject PRJNA1133156) over background i.e., Covaris fragmented genomic sequence from whole, WT planarian worms. Coordinates of the resulting MACS2-identified loci were then compared with those of the h1SMc gene annotations and overlapping loci were subtracted using BEDTools (Quinlan 2014). The remaining coordinates were then used to generate a new annotation file (.gff3; **Supplemental Material**). Transposable element annotations were generated using the Earl Grey program with default parameters (Baril *et al*. 2024)(https://github.com/TobyBaril/EarlGrey). The GO (gene ontology) enrichment analysis was done using enricher function from package clusterProfiler (version 4.10.0) (Yu *et al*. 2012; Wu *et al*. 2021). Further quantitative and statistical analysis was performed using R (version 4.3.1) and figures were created using ggplot2 (Wickham 2009).

## Results

### Optimization of an RNaseH-based ribodepletion strategy

To test the hypothesis that ribodepleted RNA-seq libraries would improve the sensitivity and diversity of S.med transcripts detected, we first needed to establish a robust ribodepletion protocol from total S.med RNA. Although there are multiple commercially-produced kits for ribodepletion from total RNA, they are designed to deplete ribosomal RNAs (rRNAs) with sequences similar to those of mammals. As the rRNA sequences of the planarian species *Schmidtea mediterranea* (S.med) are not well conserved with those of mammals, particularly the mitochondrial encoded rRNAs (**Figure 1a, Supplemental Figure S1**), most commercial kits do not work as efficiently in S.med as they do in other species. To address this issue, a previous study designed custom biotinylated probes complementary to the sequences of S.med rRNAs and showed that this strategy was highly effective for depleting rRNA from total RNA isolated from planarian stem cells (Kim *et al*. 2019b). However, both commercial kits and custom biotinylated probes are expensive options for this strategy; we therefore sought to develop a method that was similarly effective at ribodepletion but also significantly more cost-effective. We found two robust, cost-effective protocol for custom rRNA depletion and RNA-seq in bacterial and human samples (Culviner *et al*. 2020; Baldwin *et al*. 2021) and set out to optimize a protocol for S.med RNA (**Figure 1b, Supplemental Figure S2**). This strategy uses standard, untagged DNA oligos that anneal to the desired RNA sequences and then digests these RNA:DNA hybrids with RNaseH, an enzyme that specifically degrades hybrid species (Garces and Wittek 1991). The excess DNA probes are then digested with DNaseI, leaving behind a ribodepleted sample containing all remaining RNA species i.e., mRNAs, small RNAs, non-coding RNAs, etc. (**Figure 1b, c**). This RHaseH-ribodepleted (RH-ribodepleted) sample can then be used to generate an RNA-seq library (**Figure 1b**).

**Figure 1.**
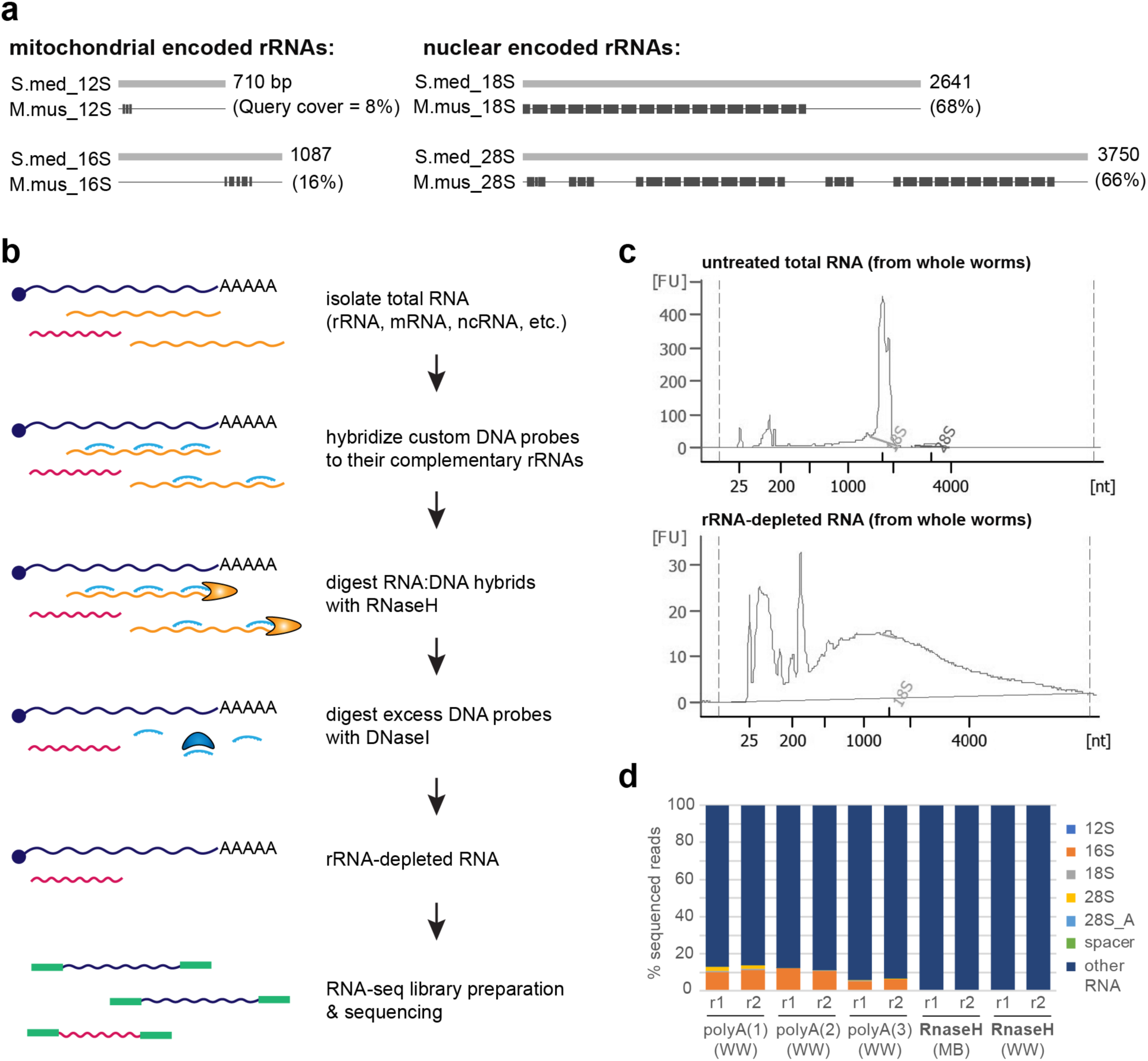
A customized, RNaseH-based ribodepletion protocol produces planarian RNA-seq libraries with consistently low rRNA reads. a) Graphic representation of the BLAST-based overlap between *Schmidtea mediterranea* (S.med) and mouse (M.mus) rRNA sequences, including both mitochondrial (12S, 16S) and nuclear-encoded (18S, 28S) rRNA species. b) Schematic of the customized, RNaseH-based method used for generating ribodepleted RNA-seq libraries from total S.med RNA. c) Bioanalyzer plots of total RNA with no rRNA depletion (top) and after ribodepletion with our optimized protocol. d) Plot showing the percentage of rRNAs detected in several published polyA-enriched RNA-seq libraries (Wong *et al*. 2022) versus multiple ribodepleted libraries prepared with the method shown in b. MB = mid-regeneration blastema tissue, WW = whole worm tissue. All libraries in d) were sequenced using paired-end technology with 2 x 150bp read lengths.

Similar to previous results produced using a biotinylated rRNA probe method and total RNA from isolated planarian stem cells (Kim *et al*. 2019b), we found that our customized S.med ribodepletion strategy generated RNA-seq libraries with consistently low (<1%) amounts of rRNA reads (**Figure 1c, d**). This is a significant reduction from the typical percentage of S.med rRNA reads detected in RNA-seq libraries generated with the more commonly used polyA-enrichment strategy (**Figure 1d**)(Kim *et al*. 2019b). To ensure that the RNAs of interest were not also degraded in this enzyme-based ribodepletion process, we optimized the conditions of RNaseH digestion by treating total RNA samples in mock reactions without DNA probes (**Supplemental Figure 2a**). We found that we could achieve similar levels of ribodepletion and limit unintended RNA degradation by reducing the RNaseH digestion time to 2.5 minutes (**Supplemental Figure 2b**). We then used this reduced time in all our library preparations.

After establishing an effective ribodepletion protocol (**Figure 1**), we used it to generate RNA-seq libraries from several different types of S.med tissues including whole worm (WW) and injury-induced regeneration blastema tissue (e.g., MB = mid-regeneration blastemas). We then sequenced the resulting libraries with paired-end sequencing technology (2 x 150bp read length) to ensure accurate read alignment and detection. Having confirmed that these libraries were efficiently rRNA depleted (**Figure 1d**), we wanted to compare the “other” RNA transcripts in these libraries (i.e., mRNA and putative noncoding RNAs) with those in polyA-enriched RNA-seq libraries. We evaluated several available RNA-seq datasets in the Gene Expression Omnibus (GEO) and download a published set of RNA-seq libraries (Wong *et al*. 2022) that are highly similar to our libraries in their starting tissue (Whole Worm, or WW), sequencing technology, depth, and percent alignment to the S.med S3h1 genome (**Table 1**) but generated using a standard polyA-enrichment strategy. We also ran a test of library complexity on both the polyA-enriched and RH-ribodepleted libraries and did not observe any notable difference (**Supplemental Figure S3**).

**Table 1.**
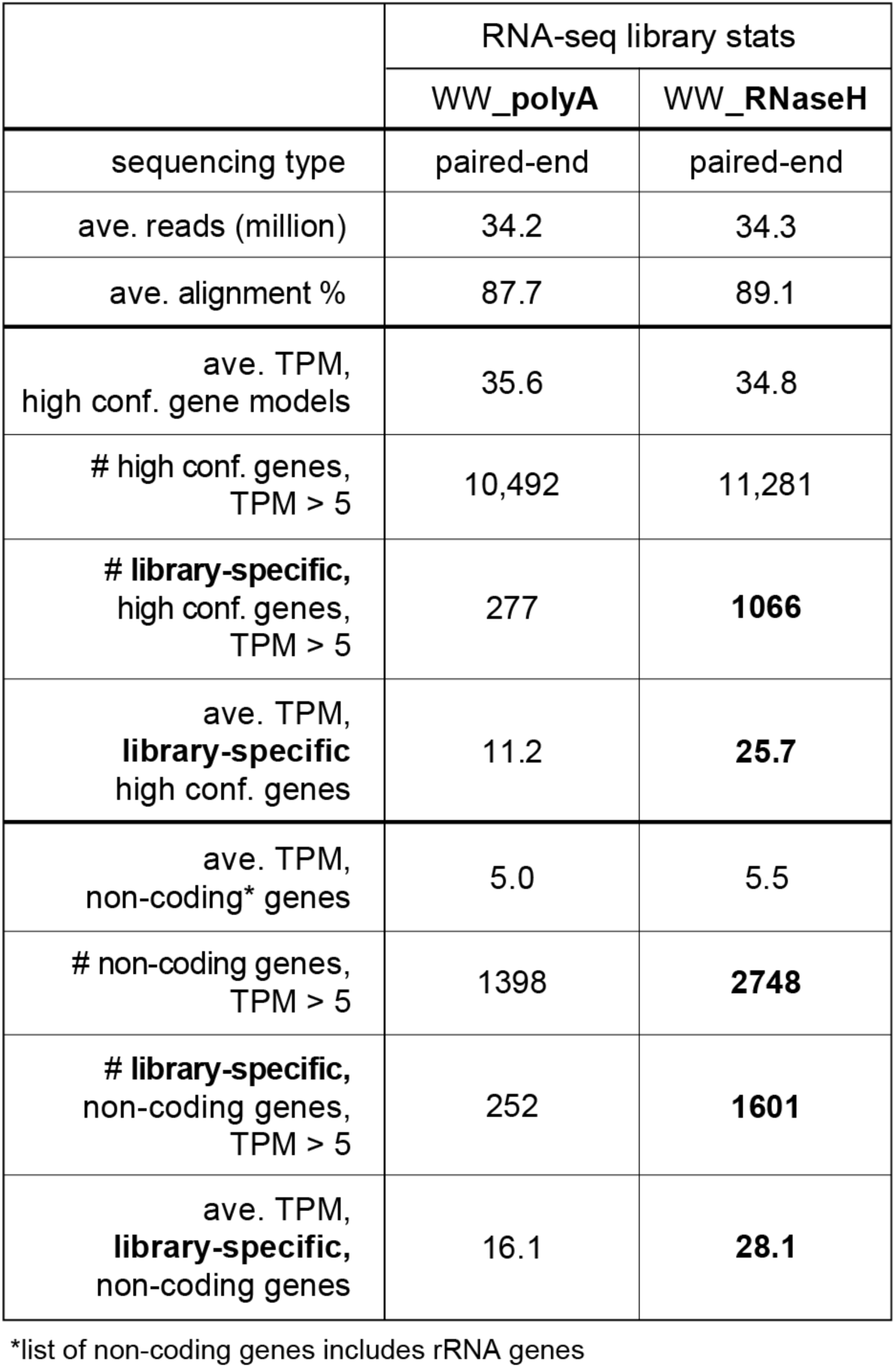
Features of the polyA-enriched and ribodepleted RNA-seq libraries used for comparison in. **Figures 2-5**.

### Comparison of transcript detection in polyA-enriched vs RH-ribodepleted RNA-seq libraries

Given their similarity in read depth and complexity (**Table 1, Supplemental Figure S3**), we compared the detection of individual gene models in these polyA-enriched RNA-seq libraries (polyA_WW) and those we generated with our customized and optimized ribodepletion strategy (RNaseH_WW; **Figure 2a**). We first looked at high confidence gene models i.e., as defined in (Ivankovic 2023) and annotated with the prefix h1SMc. Overall, these genes have a similar average expression (in Transcripts Per Million mapped reads, or TPMs) in both polyA-enriched and RH-ribodepleted libraries (**Figure 2a** and **Table 1**). However, a large subset of h1SMc genes (1,066) were only detected robustly (average TPM > 5) in the RH-ribodepleted libraries (**Table 1** and **Figure 2b**). We also found that some genes were only robustly detected (TPM > 5) in the polyA-enriched libraries, but this subset was smaller (277) and had a lower average TPM than those robustly detected in the RNaseH libraries only (**Table 1** and **Figure 2b**). We then wanted to know if there were any common biological functions among the genes that are only robustly detected in RH-ribodepleted libraries (“RNaseH-enriched” genes), as we predicted they would include specific types of transcripts; for example, it is known that those encoding core histone proteins are not polyadenylated (Ingledue *et al*. 1995; Marzluff *et al*. 2008).

**Figure 2.**
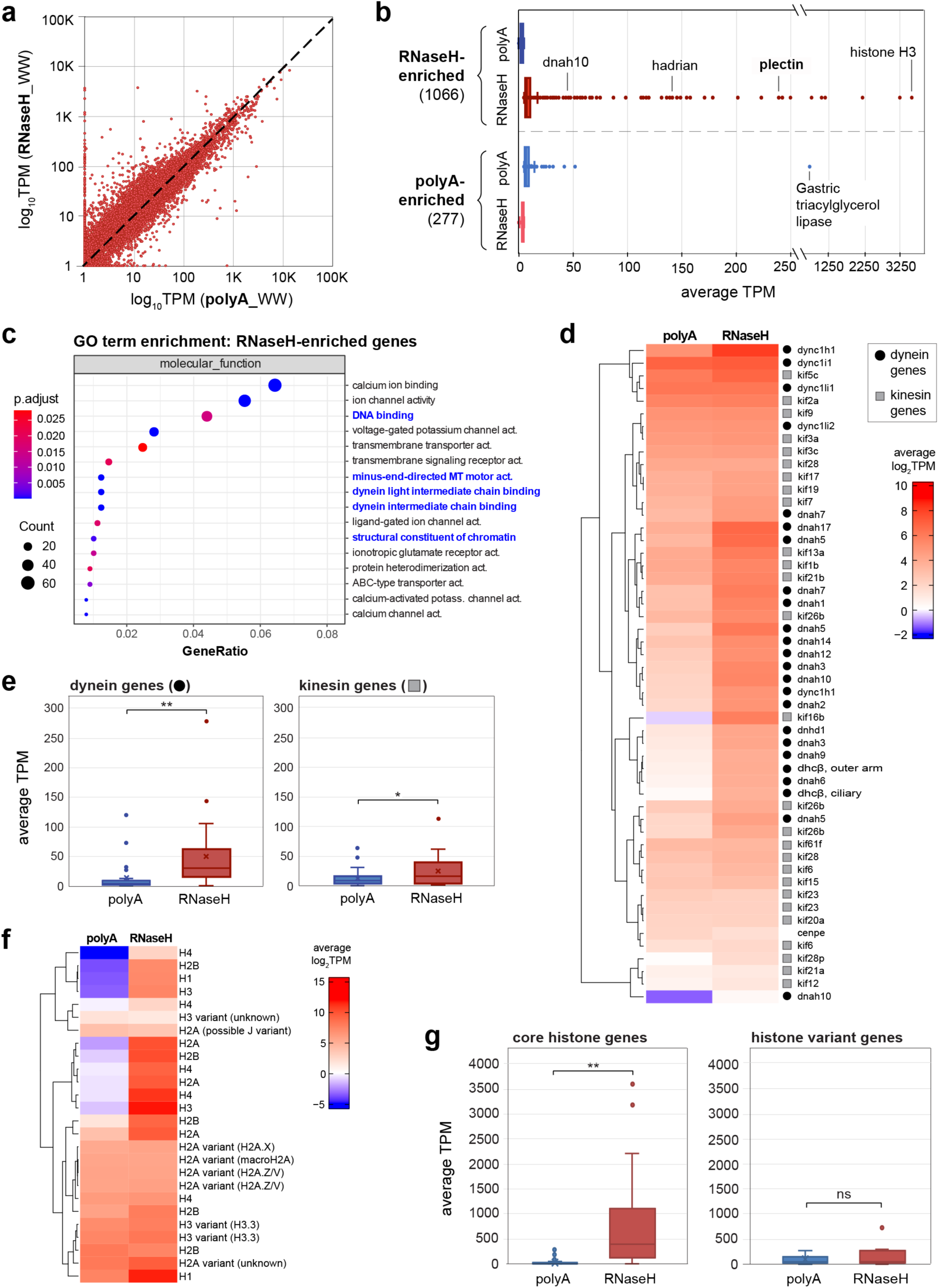
RNaseH ribodepleted RNA-seq libraries show increased detection of certain types of transcripts. a) Correlation plot comparing transcript expression (in transcripts per million mapped reads, TPM) in RNaseH-ribodepleted libraries versus polyA-enriched RNA-seq libraries; WW = whole worm tissue. b) Box plot comparing transcript expression for RNaseH-enriched genes (those with TPM > 5 in RNaseH libraries but < 5 TPM in polyA libraries) versus polyA-enriched genes (those with TPM > 5 in polyA libraries but < 5 TPM in RNaseH libraries). c) GO term analysis for RNaseH-only genes according to their molecular function. Terms in blue font are further analyze in d-g. d) Clustered heat map showing expression (log2TPM) of all gene models annotated with the GO term “microtubule-based movement”. Dynein genes are labeled with black circles, kinesin genes are labeled with gray squares. e) Box plots comparing relative expression of dynein genes (left plot) and kinesin genes (right plot) in both polyA-enriched and RnaseH-ribodepleted libraries. f) Clustered heat map showing expression (log2TPM) of all gene models annotated with the GO term “structural component of chromatin”. g) Box plots comparing relative expression of core histone genes (left plot) and variant histone genes (right plot) in both polyA-enriched and RNaseH-ribodepleted libraries. * = p-value < 0.05, ** = p-value < 0.01 (Student’s T-test).

Indeed, when we ran a Gene Ontology (GO) term enrichment analysis on the list of 1,066 RNaseH-enriched genes we found that terms such as “DNA binding” and “structural constituent of chromatin” were significantly enriched (**Figure 2c**). Unexpectedly, we also noted multiple terms associated with genes encoding microtubule motor proteins i.e., “minus end directed microtubule (MT) motor activity”, “dynein light intermediate chain binding”, and “dynein intermediate chain binding” (**Figure 1c**). However, as this GO term analysis was only done on a subset of genes meeting specific criteria (average TPM > 5 in RH-ribodepleted RNA-seq libraries and TPM < 5 in polyA-enriched libraries), we decided to perform additional analyses examining all genes annotated with these GO terms.

First, we identified all genes annotated with the GO term “microtubule-based movement” and clustered them hierarchically based on log2TPM (**Figure 2d**). As suggested by the enrichment of GO terms specific for dynein in **Figure 2c**, we observed that genes encoding conserved dynein proteins (black circles) not only clustered together but also showed a greater log2TPM in RH-ribodepleted RNA-seq libraries than polyA-enriched libraries (**Figure 2d**). To assess the significance of this observation, we averaged the detection of all conserved dynein genes (i.e., those encoding proteins annotated with the GO term “minus end directed MT motor activity” and BLAST homology to dynein proteins) and compared it with that of all conserved kinesin genes (i.e., those encoding proteins with BLAST homology to kinesins) in polyA-enriched versus RH-ribodepleted libraries (**Figure 2e**). Although both had statistically higher average TPMs in RH-ribodepleted versus polyA-enriched libraries, the difference was larger and more significant for dynein genes (p-value of dynein genes comparison < 0.01; p-value of kinesin genes comparison < 0.05).

We then identified all genes with the GO term “structural constituent of chromatin” and clustered them hierarchically based on log2TPM (**Figure 2f**). Notably, this list of histone-encoding genes includes homologs of both core, replication-dependent (RD) histones and “variant”, replication-independent (RI) histones (TALBERT AND HENIKOFF 2017). As core histones do not have polyA tails (Marzluff *et al*. 2008) but variants often do (TALBERT AND HENIKOFF 2017), we expected to see these two types of histone genes cluster distinctly. With a few exceptions, this was largely what we observed (**Figure 2f**). We then split this list of histone genes into “core” and “variant” histones based on their sequence homology and performed an independent analysis of their detection in polyA-enriched versus RH-ribodepleted libraries (**Figure 2g**). As expected, S.med gene models encoding likely “core” histones were much more robustly detected in the RH-ribodepleted libraries (**Figure 2g, left**; p-value <0.01), whereas “variant” histone genes did not show significant differences in detection between library type (**Figure 2g, right**; p-value = 0.38).

The enhanced detection of core histone gene transcripts in RH-ribodepleted RNA-seq libraries was unsurprising, as it has a clear molecular basis and has been shown in prior studies (Cui *et al*. 2010; Kim *et al*. 2019b). However, we were unsure why dynein encoding genes would be so distinctly enriched in these libraries: we could not find any reports suggesting their transcripts are not polyadenylated in other organisms and those transcripts we checked individually did have stretches of A’s at their 3’ end. However, when looking at the alignment files of the RNA-seq libraries using the Integrative Genomics Viewer (IGV) (Robinson *et al*. 2011), we noted two details: 1) dynein genes are very long compared to other genes and 2) the alignments of polyA-enriched RNA-seq libraries show strong 3’ end bias (**Figure 3a**), an issue that has been previously reported (Shi *et al*. 2021; VISCARDI AND ARRIBERE 2022). We then asked if RNaseH-enriched gene transcripts (1,066) are on average longer than those better represented in polyA libraries (277); indeed, RNaseH-enriched transcripts are significantly longer, on average, than those enriched in polyA libraries (**Figure 3b**). Moreover, when we grouped all highly conserved genes (i.e., h1SMc gene models) by transcript length, we found an increasingly significant improvement in transcript detection in RH-ribodepleted libraries when transcripts are longer than 1kb (**Figure 3c**). To address why a smaller group of genes (277) are better detected by the polyA-enrichment strategy, we asked if those transcripts have higher AT percentage (or lower GC percent) in their sequences, which could mediate oligo dT priming at regions upstream of the polyA tail. Predictably, we did find that the 277 genes more robustly detected in polyA-enriched RNA-seq libraries have a lower average GC percentage (or higher AT percentage) than either those enriched in the RNaseH libraries or the average coding (h1SMc) transcript (**Figure 3d**).

**Figure 3.**
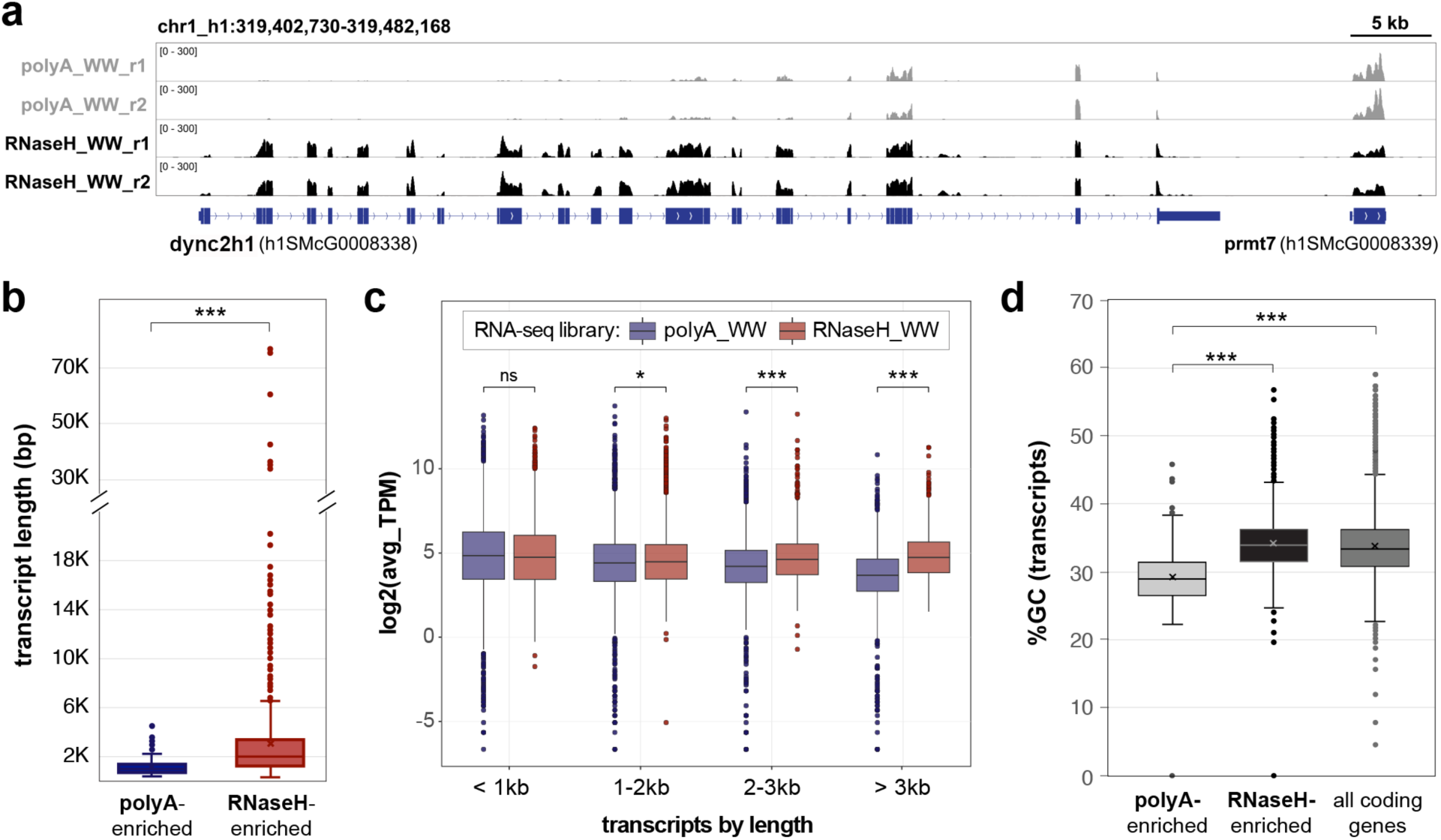
Ribodepleted RNA-seq libraries generate more complete coverage of longer transcripts. a) Representative tracks of RNA-seq data comparing polyA-enriched RNA-seq libraries and RNaseH-ribodepleted libraries at an 80kb region of chromosome 1 with two genes, cytoplasmic dynein 2 heavy chain 1 (dync2h1; h1SMcG0008338) and protein arginine methytransferase 7 (prmt7; h1SMcG0008339); WW = whole worm tissue, r = replicate. b) Box plot comparing the transcript lengths of polyA-enriched versus RNaseH-enriched genes. c) Box plot comparing the average expression (log2TPM) of transcripts grouped by length (<1 kb, 1-2 kb, 2-3 kb, and > 3kb) in polyA-enriched libraries versus RNaseH-ribodepleted libraries. d) Box plot comparing the percent of GC nucleotides in polyA-enriched, RNaseH-enriched, and all coding transcripts (i.e., those with the prefix h1SMc). * = p-value < 0.05, ** = p-value < 0.01, *** = p-value < 0.001 (Student’s T-test).

### Identifying unannotated RNAs using RH-ribodepleted RNA-seq libraries

We then asked if our RNaseH-ribodepletion strategy improved detection of putative non-coding RNAs (ncRNAs). First, we assessed if the annotated gene models with the prefix h1SMn (Ivankovic 2023) were more robustly detected in the RH-ribodepleted libraries versus polyA-enriched ones. Although these gene models are not definitively classified as ncRNAs, they do include known ncRNAs (e.g., ribosomal RNAs) and likely include many others given that they lack or have very short open reading frames (ORFs). As with the high confidence gene models (prefix h1SMc), the h1SMn models have similar overall average TPMs in both polyA-enriched and RH-ribodepleted libraries. However, a large subset of them (1,601) were only detected robustly (average TPM > 5) in the RH-ribodepleted libraries (**Table 1**). We also identified h1SMn genes that are better detected in the polyA-enriched libraries, but this subset was smaller (252) and had a lower average TPM than those better detected in the RNaseH libraries (**Table 1** and **Supplemental Figure S4**).

In addition to showing that our RNaseH-ribodepletion method offers improved detection of annotated gene models, we predicted that this strategy would detect many putative ncRNA transcripts that are not represented in these models as the current annotations are largely based on RNA-seq data generated using polyA-enrichment strategies (Ivankovic 2023) and many ncRNAs are not polyadenylated (Barrett *et al*. 2021; Limberis *et al*. 2023). In support of this hypothesis, we observed many unannotated loci with substantial and consistent signal in RH-ribodepleted libraries but not polyA-enriched libraries when examining RNA-seq alignments using IGV (**Figure 4a**, orange shading). To assess if this signal was significant and to find similar loci across the entire S.med genome, we designed a bioinformatics pipeline to identify regions with enriched RNA-seq signal over background; we then subtracted those that overlap with existing high confidence gene models (h1SMcG; **Figure 4b**). Using this pipeline, we identified over 14K genomic loci that were ≥ 200bp (**Table 2** and **Figure 4c, d**). These loci are scattered across the genome on all four chromosomes and in both gene dense and intergenic regions (**Figure 4c**). Because we only subtracted high-confidence gene models (h1SMc), we found that over one third of these loci overlapped with h1SMn genes models (**Table 2**). To further verify that this signal represents active transcription at these loci, we designed primers to several identified regions (marked by orange shading in **Figure 4a, c**) and performed PCR with cDNA synthesized from an independent sample of total S.med RNA (**Figure 4e**). We detected bands at the predicted size for all primer pairs mixed with cDNA, but not in the -RT control reactions (**Figure 4e**). Although we did also test if cDNA synthesis with random hexamers (R) versus oligo dT primer (O) was required to detect these transcripts, we only noted small or no differences in detection (**Figure 4e**). However, given we used gene specific primers and standard PCR conditions with 30 cycles of amplification, our detection of these transcripts is both more sensitive and less quantitative than other expression assays.

**Figure 4.**
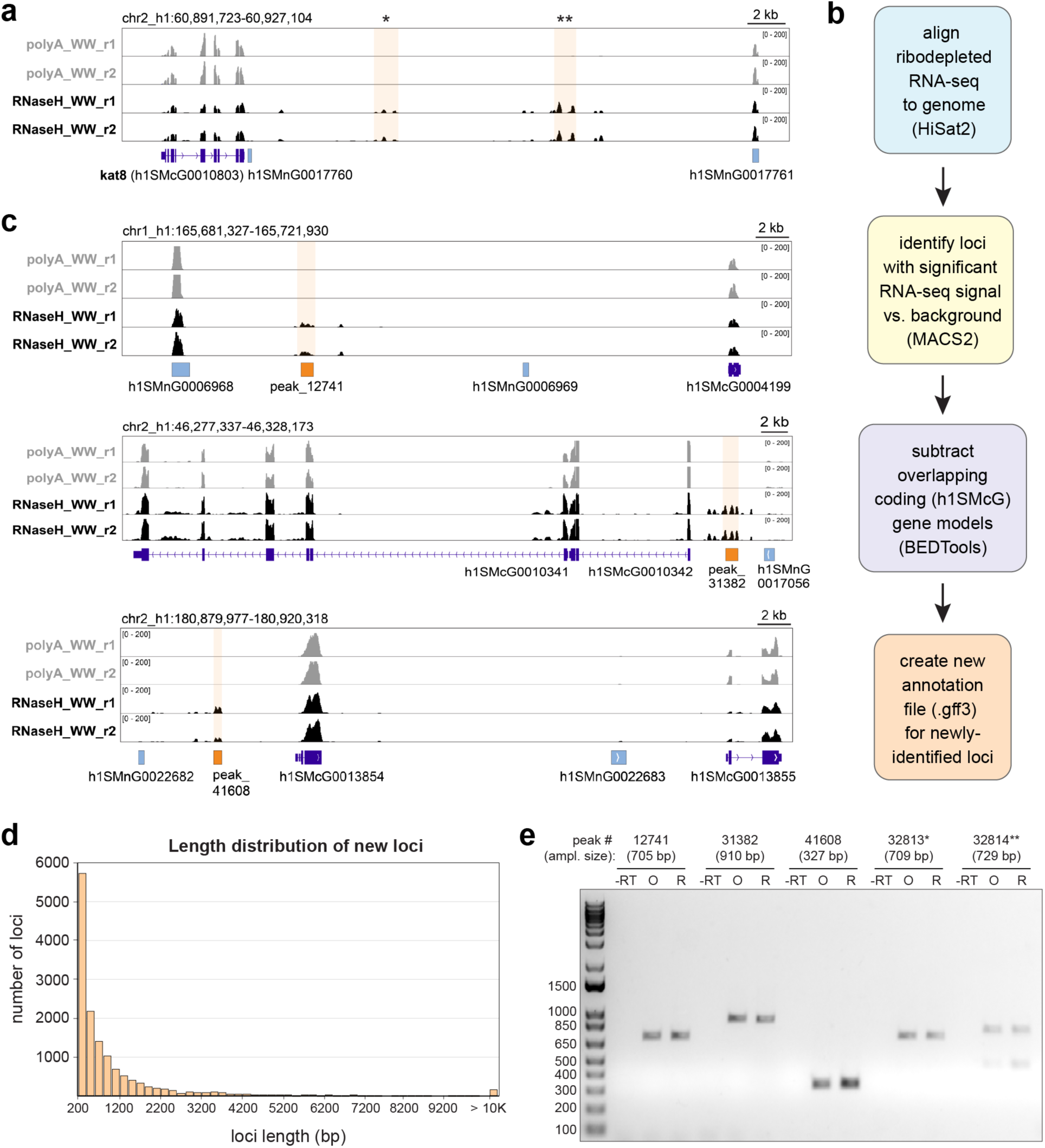
Ribodepleted RNA-seq libraries contain reads that map to regions of the S3h1 genome where there are no annotated gene models. a) Representative track of RNA-seq data from polyA-enriched and RNaseH-ribodepleted libraries at a region of the S3h1 genome where there are detectable peaks of aligned RNA-seq reads that do not correspond to an annotated gene model (orange shading); WW = whole worm tissue, r = replicate. b) Schematic describing the pipeline developed to identify loci with significant reads but no high confidence (h1SMcG) gene model. c) Representative tracks of RNA-seq data from polyA-enriched and RNaseH-ribodepleted libraries at three separate regions of the S3h1 genome with loci (orange shading) detected by the pipeline outlined in b. d) Histogram showing the distribution of lengths of the loci identified. e) Gel showing products of PCR reactions with primers designed to each RNaseH-library-specific locus highlighted in a and c. Shaded region in a) marked by * = peak_32813, ** = peak_32814. “-RT”, “O”, and “R” indicate type of cDNA template used in PCR reaction; -RT = no reverse transcriptase control, O = cDNA synthesized with oligo dT priming, R = cDNA synthesized with random hexamer priming.

**Table 2.**
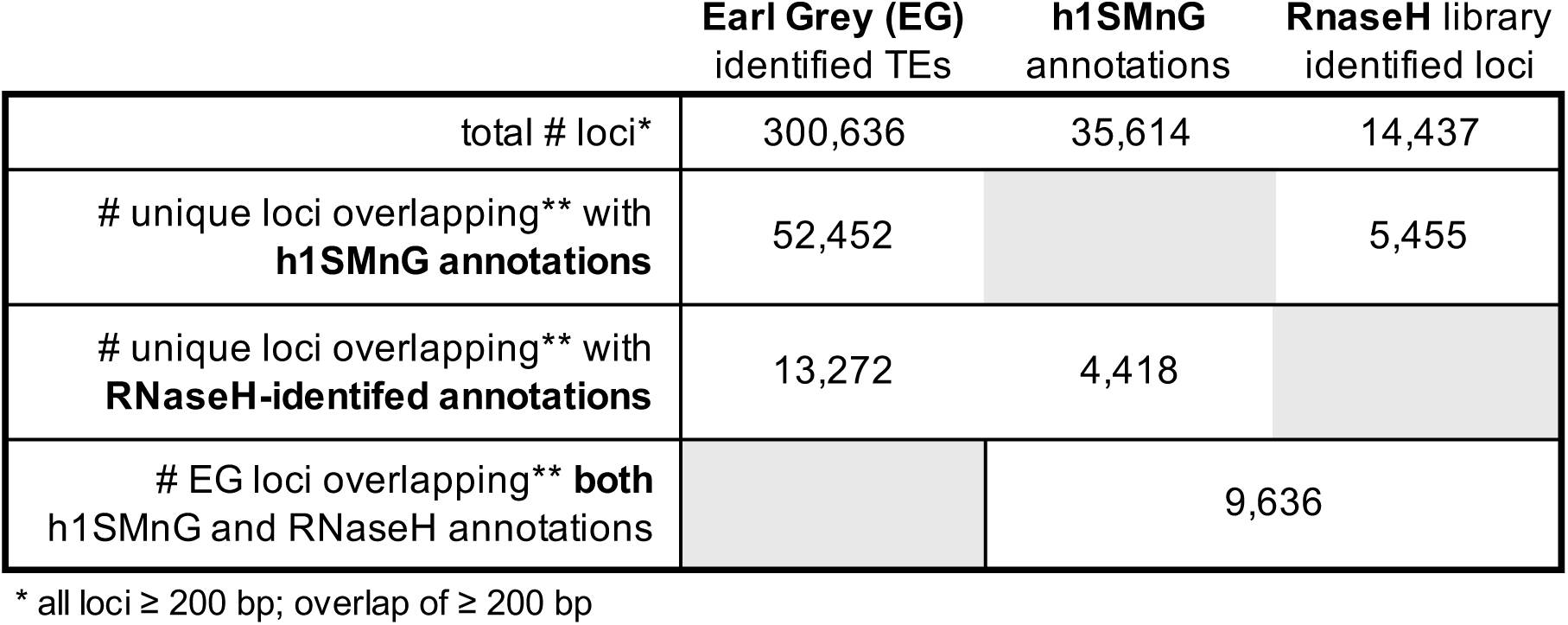
Summary of putative ncRNA loci identified using multiple resources and bioinformatic tools.

After identifying thousands of new loci, we wanted to understand what these transcripts may represent. One likely possibility is that they are transcribed Transposable Elements (TEs). Although TEs were historically thought to be constitutively silenced in most genomes, recent evidence shows that they are often expressed, that their expression is regulated, and that this regulated expression plays essential roles in cell biology (SIMO-RIUDALBAS *et al*. 2022; Oomen and torres-padilla 2024; Wang *et al*. 2024). To address whether some or all of our newly identified loci are TEs, we first analyzed the S3h1 genome using a recently published TE classification program called Earl Grey (Baril *et al*. 2024). As expected based on previous examinations of the S.med genome, this program identified sizeable percentages of the S.med genome as likely TEs (**Figure 5a**). We then compared these genome-wide Earl Grey (EG) classifications with both the h1SMn gene models and the new, putative ncRNA loci we identified using our RH-ribodepleted libraries (RHnc). This comparison showed that our newly identified loci represent many putative DNA and LTR (Long Terminal Repeat) elements (**Figure 5b, c**). Moreover, we again verified with PCR that these likely TE elements are transcribed into RNA (**Figure 5d**).

**Figure 5.**
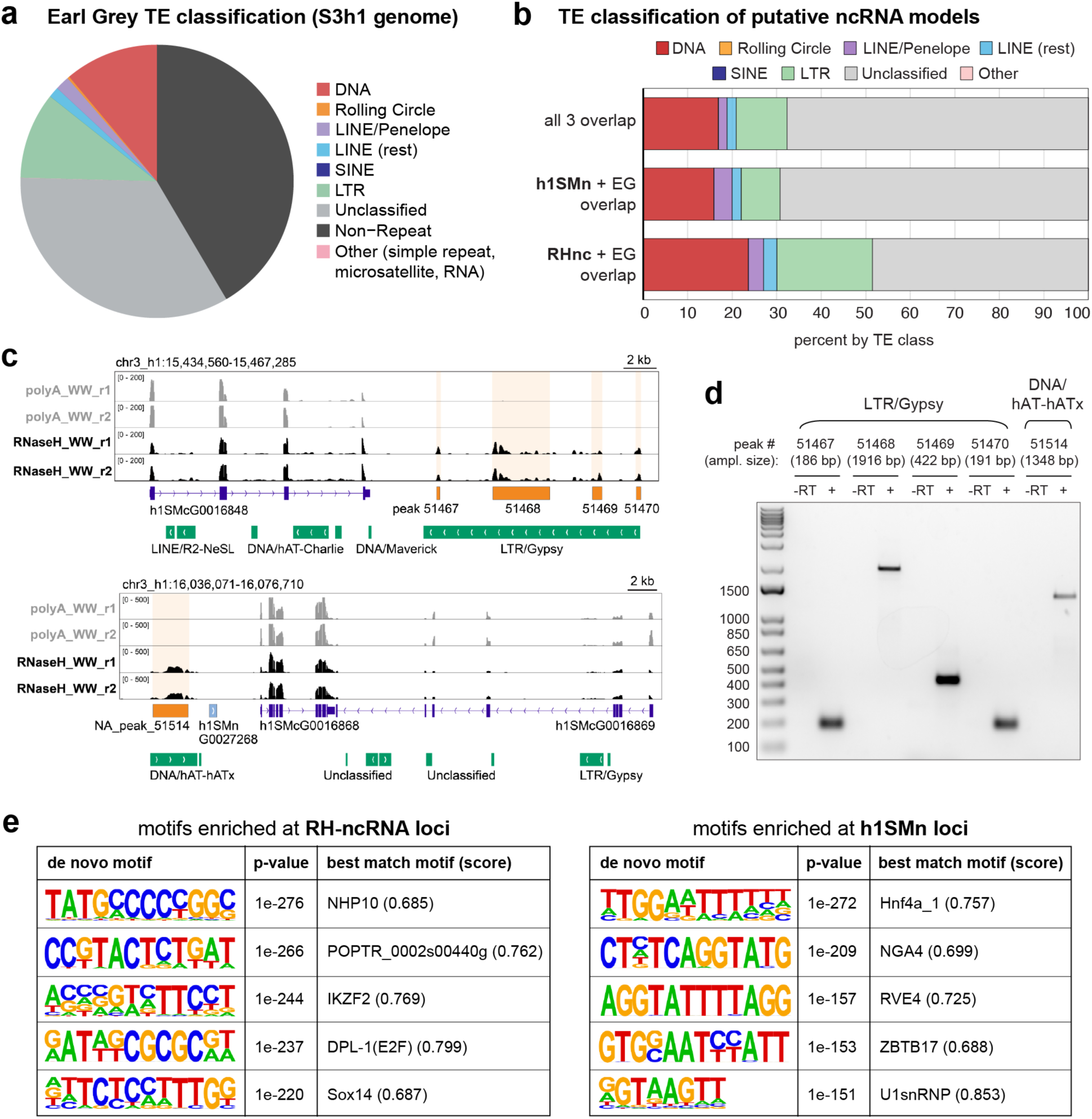
RNaseH ribodepleted libraries detect both novel loci as well as many that overlap with independently generated annotations. a) Pie chart summarizing the types of Transposable Elements (TEs) detected in the S3h1 genome by the Earl Grey program (Baril *et al*. 2024). b) Plot comparing the types of Earl Grey-detected TEs in the putative non-coding RNAs gene models in the S3h1 annotations (i.e., those starting with h1SMn), the new putative non-coding RNA loci detected using our RNaseH-ribodepleted libraries (RHnc), and those found in both (all three overlap). Overlap = > 200bp overlap between annotations. c) Representative tracks of RNA-seq data from polyA-enriched and RNaseH-ribodepleted libraries at two separate regions of the S3h1 genome with loci (orange shading) detected by both the pipeline outlined in Figure 4b and the Earl Grey TE detection program. d) Gel showing products of PCR reactions with primers designed to each RNaseH-library-specific locus highlighted in c. “-RT” and “+” indicate type of cDNA template used in PCR reaction; -RT = no reverse transcriptase control, + = cDNA synthesized with random hexamer priming. e) de novo motif analysis (HOMER) of potential DNA binding motifs enriched in the loci uniquely identified by the pipeline outlined in Figure 4b versus loci uniquely identified by the h1SMn gene annotations.

Many recent studies have focused on the finding that some TEs, particularly ERVs (Endogenous Retroviral Elements) in the LTR class, encode numerous transcription factor (TF) binding motifs (GARRIGUES AND PASQUINELLI 2022; Sekine *et al*. 2023; Hossain *et al*. 2024). To assess whether our newly identified loci also encode such motifs, we performed de novo motif analysis using HOMER (Heinz *et al*. 2010). Notably, this analysis identified many motifs in the set of RNaseH-library-specific loci that were both significantly enriched and distinct from those identified in the h1SMn gene model sequences (**Figure 5e**). Although this analysis cannot determine the relevance of the identified motifs, it does suggest that our newly identified loci have distinct characteristics.

## Discussion

Planarian flatworms are a model organism with fascinating biology and enormous potential to uncover pathways, mechanisms, and molecules that inform questions about aging, stem cells, regeneration, and even cancer. Although scientists have been studying these animals for hundreds of years (ELLIOTT AND SANCHEZ ALVARADO 2013) and using forward genetic tools to identify important planarian genes for decades (SANCHEZ ALVARADO AND NEWMARK 1999; NEWMARK AND SANCHEZ ALVARADO 2002; Reddien *et al*. 2005), the field is now in a new phase of tool building and transformation. In the last few years, new chromosome level genomes have been assembled (Guo *et al*. 2022; Ivankovic 2023), dozens of transcriptomes for many different species have been constructed (Almazan *et al*. 2018; Duncan *et al*. 2022; VILA-FARRE *et al*. 2023), and methods for assaying the chromatin state and dynamics have been optimized (Duncan *et al*. 2015; Ivankovic 2023; PASCUAL-CARRERAS *et al*. 2023; Poulet *et al*. 2023; Verma *et al*. 2024) to better understand the context in which these genetic blueprints exist in planarian cells.

In this study, we aimed to contribute to this exciting phase of growth in planarian genomics and transcriptomics by developing an approach that captures the depth and complexity of planarian RNA transcripts. Different from other studies that use RNA-seq to measure differential expression between samples, here we provide an optimized method, a wealth of resources, and a tested bioinformatics pipeline that can all serve as a foundation for future experiments.

Moreover, we find that our method offers specific advantages over more commonly used approaches in our field: we observe more robust detection of particular types of transcripts (e.g., core histones), improved coverage of longer transcripts, and the detection of transcripts not currently annotated on the planarian genome (putative ncRNAs, eRNAs, expressed TEs, etc.). Further examination of this data points to opportunities for exciting future studies, including the possibility of using our comparisons of histone transcript detection in different types of RNA-seq libraries to identify new histone variants with uncharacterized functions. Although it is arguable that we are not likely to identify many new protein coding genes with our data and pipeline given the depth of existing transcriptomic data in the field, we note that all histone H4 encoding transcripts are currently included in the “non high confidence” gene annotations (with prefix h1SMn) on the S3h1 genome, likely due to their ORF length cutoff (Ivankovic 2023). It is therefore possible that our newly identified loci include transcripts that encode small proteins or peptides with important biological functions.

Importantly, we acknowledge that we are mapping RNA-seq reads generated from the asexual S.med biotype to a genome assembled from sexual S.med sequence. Although this is common practice in our field because a asexual S.med genome assembly of comparable quality is not available, it is an issue we must consider in drawing any conclusions or designing any experiments based on these findings. In the future, when a high-quality, chromosome-level S.med asexual genome assembly is available, we and/or others can easily map these data again and compare the outputs of our pipeline. However, we also posit that the new loci identified here may represent those regions that are highly conserved between the two biotypes, as our RNA-seq reads map with high alignment rates (88-90%) to the sexual S3h1 genome using standard stringency requirements. Despite this caveat, our data and analyses contribute valuable resources for our field, especially for those with interest in ncRNAs and TEs. We have not only identified thousands of transcripts that likely represent many different RNA species (e.g., enhancer RNAs, long noncoding RNAs, TEs), but also mapped them to the genomic loci from which they are transcribed. Given the strong interest into the role of TE loci during early mammalian development (Fueyo *et al*. 2022; Oomen and torres-padilla 2024; Wang *et al*. 2024) and the evolution of gene regulatory regions (Sundaram and wysocka 2020; SENFT AND MACFARLAN 2021; Sekine *et al*. 2023; Hossain *et al*. 2024), we are excited to follow up on our observation that many of the transcripts we identified using RNaseH-depleted RNA-seq libraries are distinct from the current gene annotations and their genomic loci have distinctly different TF binding motifs. In future studies, we will investigate when these loci are activated and/or silenced, the mechanisms that accomplish this dynamic regulation, and the downstream effects of their transcription.

## Data Availability

The Bioproject (PRJNA1133156) for all new genomic sequencing data reported here is available at: https://dataview.ncbi.nlm.nih.gov/object/PRJNA1133156?reviewer=6bd87mgnoje2p40uknem <u>mialup</u>. The polyA-enriched RNA-seq libraries used for comparison can be found using GEO accession GSE174228 (three control replicate libraries).

## Acknowledgements

The authors would like to thank the DNA Services Lab in the Roy J. Carver Biotechnology Center at the University of Illinois Urbana Champaign for their expert advice and professionalism in providing sequencing services. We would also like to thank Laura Banaszynski and Jeremiah Smith for their valuable feedback on the manuscript.

## Funding

This work was funded by NIGMS grants 5R35GM142679 (E.M.D.)

## Conflict of Interest Statement

The authors state they have no conflicts of interest.

## SUPPLEMENTAL MATERIAL

Supplemental information includes four figures, three tables, and two genome annotation files:

Supplemental Figure S1. (alignments of mouse and S.med 12S and 16S rRNA sequences)

Supplemental Figure S2. (optimization of RNaseH digestion time)

Supplemental Figure S3. (comparison of complexity in polyA vs RNaseH-generated libraries)

Supplemental Figure S4. (comparison of putative noncoding h1SMn transcript detection in polyA

Supplemental Table S1 (sequences of DNA oligos used for S.med rRNA depletion)

Supplemental Table S2 (TPM data for h1SMc/h1SMn genes, polyA and RNaseH WW libraries)

Supplemental Table S3 (genome coordinates for loci identified by pipeline in Figure 4b)

Supplemental File 1 (BED file of Earl Grey output for the S3h1 genome)

Supplemental File 2 (.gff3 file of new gene annotations identified using the pipeline in Figure 4b)

## Notes

### Competing Interest Statement

The authors have declared no competing interest.

