## Supplemental Figures for "RNaseH-based ribodepletion of total planarian RNA improves detection of longer and non-polyadenylated transcripts"

### Barai P. et al, Supplemental Figures

**a** **12S mitochondrial rRNA alignment (59.2% identity)**

```
S.med --------------------------TAAGTACAATTTCTATTATACTTAC-------CA    27
M.mus AAAAGGTTTGGTCCTGGCCTTATAATTAATTAGAGGTAAAATTACACATGCAAACCTCCA    60
                                *** ** *  *   **** ** * *       **

S.med TGTTACGACTTAACTCCTCTTAAAGAAC----GGAGATTGACGGGCGGTGTGTACTAAGT    83
M.mus TAGACCGGTGTAAAATCCCTTAAACATTTACTTAAAATTTAAGGAGA--GGGTATCAAGC    118
      *    **   ***   * ****** *        * *** * **     * ***  ***

S.med CC-----------------------------------------------------TAGAA    90
M.mus ACATTAAAATAGCTTAAGACACCTTGCCTAGCCACACCCCCACGGGACTCAGCAGTGATA    178
       *                                                     *   *

S.med AAGATTC---AA-AAAAGCCAATTCATCTAAATT-TACTATTTAGTCCTGAT--------    137
M.mus AATATTAAGCAATAAACGAAAGTTTGACTAAGTTATACCTCTTAGGGTTGGTAAATTTCG    238
      ** ***    ** *** *  * **   **** ** ***   ****   ** *

S.med ------------------------CAAC----ACCAATTATTTC------AAAAGGTATC    163
M.mus TGCCAGCCACCGCGGTCATACGATTAACCCAAACTAATTATCTTCGGCGTAAAACGTGTC    298
                               ***    ** ****** *       **** ** **

S.med TAACC----------AAT-GTAGTGAAAGCATAATCTAAGATAAGCAG-CACAATGATTT    211
M.mus AACTATAAATAAATAAATAGAATTGAAATCCAA--CTTATATGTGAAAATTCATTGTTAG    356
       *             *** * * ***** *  *  ** * **  * *    ** ** *

S.med ---------GCATTAAAGAAAGCATAACTACAAGGCCATTCACATATGTACAACCGTACA    262
M.mus GACCTAAACTCAATAACGAAAGTAATTC--------TAGTCATT--TATAAT-----ACA    401
                ** *** ***** *   *         * ***    * **       ***

S.med CTAACCATAATCTTAGCGAGATATTG--AGCGGATACTC---TTTTCTAAACCCACACTC    317
M.mus CGA-----CAGCTAAGACCCAAACTGGGATTAGATACCCCACTATGCTTAGC-CATAA-A    454
      * *      * ** **    * * **  *   ***** *   * * ** * * ** *

S.med CCCAAATAATTAGAACAACTGCCAAGTCTTTTCATTTTGATGAGGAAATAAAATAAACAA    377
M.mus CCTAAATAATTAAATT---TAACAAAACTA-----TTTGCCAGAGAACTA--CTAGCCAT    504
      ** ********* *     *  ***  **      ****     *** **   **  **

S.med CCCTTATAAATAAAGCCAG---------ATTCATGTCTAA--AAAGGAAG------TGAA    420
M.mus AGCTTAAAACTCAAAGGACTTGGCGGTACTTTATATCCATCTAGAGGAGCCTGTTCTATA    564
        **** ** * **   *           ** ** ** *   * ****        *  *

S.med ACCAATAAAAGT--------ATTAAAATCTACCACTAGT----CATACATAAAGTC-TCT    467
M.mus ATCGATAAACCCCGCTCTACCTCACCATCTCTTGCTAATTCAGCCTATATACCGCCATCT    624
      * * *****            * *  ****    *** *    * ** ***  * * ***

S.med TATTGAAA-CTTAAC-------CAATTTAAACCAAATAATAAACTAACATAGC------A    513
M.mus TCAGCAAACCCTAAAAAGGTATTAAAGTAAGCAAAAGAAT---CAAACATAAAAACGTTA    681
      *    *** * ***         **  *** * *** ***   * ******        *

S.med ATACAAAGTGTAACCGCAGAAGCTGGCAC---------TCATTTGATCTCCTAATAGCAT    564
M.mus GGTCAAGGTGTAGCCAATGAAATGGGAAGAAATGGGCTACATTT--TCTTATAAAAGAAC    739
         *** ***** **   ***   ** *           *****  ***  *** ** *

S.med ATACCGACTCAAGCAATCAATTAACATTAAATTTTAAAATATACACGCCTAATAAAACCA    624
M.mus ATTACTATA-------------------CCCTTTATGAAACTAAAGGACTAAGGAG----    776
      **  * *                        ***   **  ** * * ****  *

S.med TTAAATAAAATTAATAGTAAACAACCACCAATAAACTATAGCGTTTAACAGCA-------    677
M.mus -------GATTTAGTAGTAAATTAAGA--------ATAGAGAGCTTAATTGAATTGAGCA    821
              * *** *******  *  *         ** ** * ****  * *

S.med ----------------TGCCAG--------------TTTAAAAAGCCAAGACAAAACTA    706
M.mus ATGAAGTACGCACACACCGCCCGTCACCCTCCTCAAATTAAATTAAACTTAACATAATTA    881
                        *** *              ** **  *  *   *** ** **

S.med TATT---- 710
M.mus ATTTCTAG 889
        **
```

**b** **16S mitochondrial rRNA alignment (50.2% identity)**

```
S.med ------------------------------------------------------------    0
M.mus ACTAATCCTAGCCCTAGCCCTACACAAATATAATTATACTATTATATAAATCAAAACATT    60


S.med ------------------------------------------------------------    0
M.mus TATCCTACTAAAAGTATTGGAGAAAGAAATTCGTACATCTAGGAGCTATAGAACTAGTAC    120


S.med ------------------------------------------------------------    0
M.mus CGCAAGGGAAAGATGAAAGACTAATTAAAAGTAAGAACAAGCAAAGATTAAACCTTGTAC    180


S.med ------------------------------------------------------------    0
M.mus CTTTTGCATAATGAACTAACTAGAAAACTTCTAACTAAAAGAATTACAGCTAGAAACCCC    240


S.med ------------------ATAATATATTTTTATTATTTCT-------ATTTT-CCGAATA    34
M.mus GAAACCAAACGAGCTACCTAAAAACAATTTTATGAATCAACTCGTCTATGTGGCAAAATA    300
                          ** * * ****** * *          ** *  *  ****

S.med GATGATCCTTGTTTGGATACTAGAGTATTGCATTG-----TTAGTTGT------------    77
M.mus GTGAGAAGATTTTTAGGTAGAGGTGAAAAGCCTAACGAGCTTGGTGATAGCTGGTTACCC    360
      *        * *** * **   * * *  ** *       ** **  *

S.med -------------------AATATTTTCCTTTTTGTATTACGGTTCGTAGGTATTTTTAT    118
M.mus AAAAAATGAATTTAAGTTCAATTTTAAACTTGCTAAAAAAACAACAAAATCAAAAAGTAA    420
                         *** **   ***  *  *  *        *   *    **

S.med GTTTTTATTCTCTCGAATGGAATATGATATATCTTTTGTGTG-----TTTTGTTTTATTT    173
M.mus GTTTAGATTATAGCCAAAAGAGG--GACAGCTCTTCTGGAACGGAAAAAACCTTTAATAG    478
      ****  *** *  * **  **    ** *  **** **              *** **

S.med TCTTTGGTTAACAT-------------------------------TAGAATTACATT----    199
M.mus TGAATAATTAACAAAACAGCTTTTAACCATTGTAGGCCTAAAAGCAGCCACCAATAAAGA    538
      *   *  ******                                **      **

S.med ---GATTGGTAGTAGTGATATGTTTTCGG---ATTTCTTTATCT---AGTTTTTAG----    246
M.mus AAGCGTTCAAGCTCAACATAAAATTTCAATTAATTCCATAATTTACACCAACTTCCTAAA    598
           **     *    ***   ****     *** * * ** *        **

S.med -TTGTCTTTAGGTTAAGCTTTTGTTTTA---------------------------CTTGT    278
M.mus CTTAAAATTGGGTTAATCTATAACTTTATAGATGCAACACTGTTAGTATGAGTAACAAGA    658
       **    ** ****** ** *   ****                           *  *

S.med TTTTTTGTCCTATTTGTTTA-AGTTATTCTTAATAGTATTTCTTTTAATCGTTAAAGATT    337
M.mus ATTCCAATTCTCCAGGCATACGCGTATAACAACTCGGATAACCATTGTTAGTTAATCAGA    718
       **    * **    *  **    ***    * * * **  *  **  * *****  *

S.med TGTTA----GTCTTCTGTTTGTTTATATGGTTTTTAGTAAT--TCTTTTTAG-CAATTAC    390
M.mus CTATAGGCAATAATCACACTATAAATAATCCACCTATAACTTCTCTGTTAACCCAACACC    778
         **     *  **    * *  ***       **  * *  *** ** *  ***   *

S.med GTAATTAA---------TATGAATTATGCTATATATA-----ACTCAATGCGTCTACAAC    436
M.mus GGAATGCCTAAAGGAAAGATCCAAAAAGATAAAAGGAACTCGGCAAACAAGAACCCCGCC    838
      * ***             **  *  * * ** *   *      *  *      *  *  *

S.med TGTTTTCTAAAAACATTTCATTTTTTGTAAAAT---------GTAGTCCCT--GCTCACT    485
M.mus TGTTTACCAAAAACATCACCTCTAGCATTACAAGTATTAGAGGCACTGCCTGCCCAGTGA    898
      ***** * ********  * * *    * * *          * * * ***   *

S.med GATAAGTTAAATAGCTGCAGTACTTTGACTGTACGAAGGTAGCATAATTACTTGTCTATT    545
M.mus CTAAAGTTTAACGGCCGCGGTATCCTGACCGTGCAAAGGTAGCATAATCACTTGTTCCTT    958
         ***** **  ** ** ***   **** ** * ************* ******   **

S.med AATTCTAGAATTGTTTGAATGGGTTTATTGATAGATAGCAAGTTTTATTTAGTCTTGTTT    605
M.mus AATTAGGGACTAGCATGAACGGCTAAACGAGGGTCCAACT--GTCTCTTATCTTTAATCA    1016
      ****   ** * *  **** ** *  *         * *    * * **   * *  *

S.med TTTAATTTATACTTTCTGTAAAGATACAGTTTGATATTTCAAGGACGAAAAGACCCTATA    665
M.mus GTGAAATTGACCTTTCAGTGAAGAGGCTGAAATATAATAATAAGACGAGAAGACCCTATG    1076
       * ** **   ***** ** ****  * *    *** *   * ***** **********

S.med GAGTTTTTAACTTAGTGGTGTTTCTTACTTTAGTTATTTGTTGGGGTAACGGTATTTCTT    725
M.mus GAGCTTAAATTATATAACTT--ATCTATTTAATTTATTAAACCTAATGGCCCAAAAACTA    1134
      *** **  *   **    *      ** ** * *****        *  *   *   **

S.med TTGAGTACTTATTTAT--------------------------------------------    741
M.mus TAGTATAAGTTTGAAATTTCGGTTGGGGTGACCTCGGAGAATAAAAAATCCTCCGAATGA    1194
      * *  **  * *  *

S.med ------------------------------------------------------------    741
M.mus TTATAACCTAGACTTACAAGTCAAAGTAAAATCAACATATCTTATTGACCCAGATATATT    1254


S.med ------TACATTATGAACTTCCTTAGGGATAACAGGGATATAGAATCTTGGAGGACATAT    795
M.mus TTGATCAACGGACCAAGTTACCCTAGGGATAACAGCGCAATCCTATTTAAGAGTTCATAT    1314
             **      *  * ** ************ *  **   ** *  ***  *****

S.med CGAAGATTTTG-TTTTCTACCTCGATGTTGAATTGTAGTTAAAGATAGGTGTAGAGGC-C    853
M.mus CGACAATTAGGGTTTACGACCTCGATGTTGGATCAGGACATCCCAATGGTGTAGAAGCTA    1374
      ***  ***  * *** * ************ **           *  ******** **

S.med TTTACTTTTAGTCTGTTCGACTAATAATACTATTCGTGATTTGAGTTTAGACCGATGTGA    913
M.mus TTAATGGTTCGTTTGTTCAACGATTAAAGTCCTACGTGATCTGAGTTCAGACCGGAGCAA    1434
      ** *   ** ** ***** ** * ***     * ****** ****** ******  *  *

S.med ATCAGGTTGGTTTTTATCCTGAATCACTGTCCATTGTACGAAAGGAATTGGATTGATATT    973
M.mus TCCAGGTCGGTTTCTATCTATTTACGATTTCTCCCAGTACGAAAGGACAAGAGAAATAGA    1494
        ***** ***** ****      *  * **          ** * *   **   ***

S.med GTTAGATAAAAAAAAAAAAAAAAAATATTGTGTTGACCTTTCTTTCGAGGATTTTTTTTT    1033
M.mus GCCACCTTACAAAT--------------------AAGCGCTCTCAACTTAATTTATGAAT    1534
      *  *  * * ***                      * *  ***       **** *   *

S.med TTTGTTGTTTTTAGCTTTTGTGTTTCATGTTGTTTCGACGGCATTTTTTTATGT    1087
M.mus AAAATCTAAATAAAATATATACGTACACCCTCTAACCTAGAGAAGGTT------    1582
      *     * *  * *     * **   * *  *   *  *   **
```

**Supplemental Figure S1.** Alignment of S.med and mouse 12S (left) and 16S (right) rRNA sequences

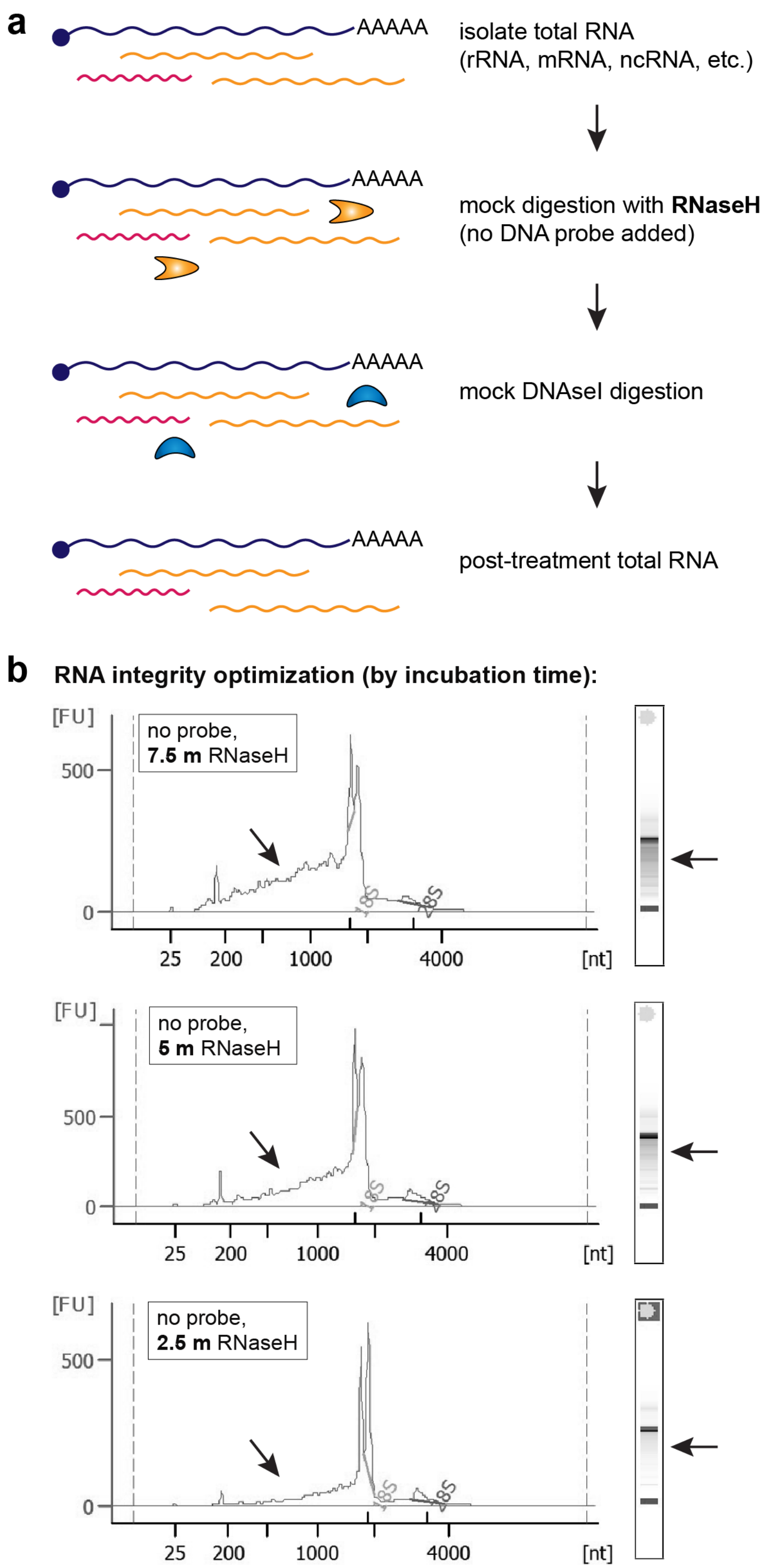


**Supplemental Figure S2.** a) Schematic of strategy for assessing secondary degradation of S.med RNA during RNaseH and DNase I digestion. b) optimization of RNaseH digestion time to reduce secondary RNA degradation. Arrows indicate RNA degradation products, which are lowest with 2.5 minutes RNaseH digestion.

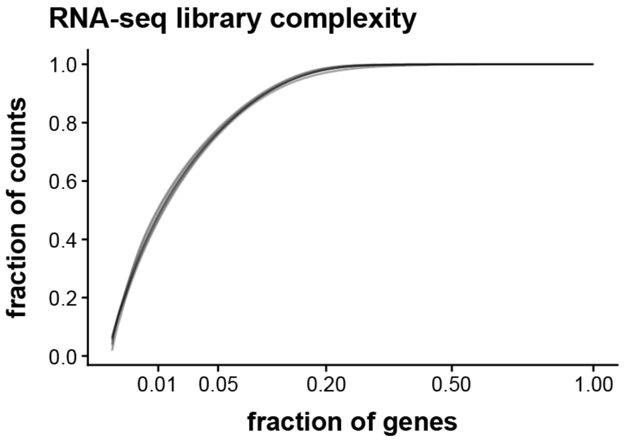


**Supplemental Figure S3.** Plot comparing complexity of all polyA-enriched libraries (grey lines) and RNaseH-ribodepleted libraries (black lines) analyzed in this study.

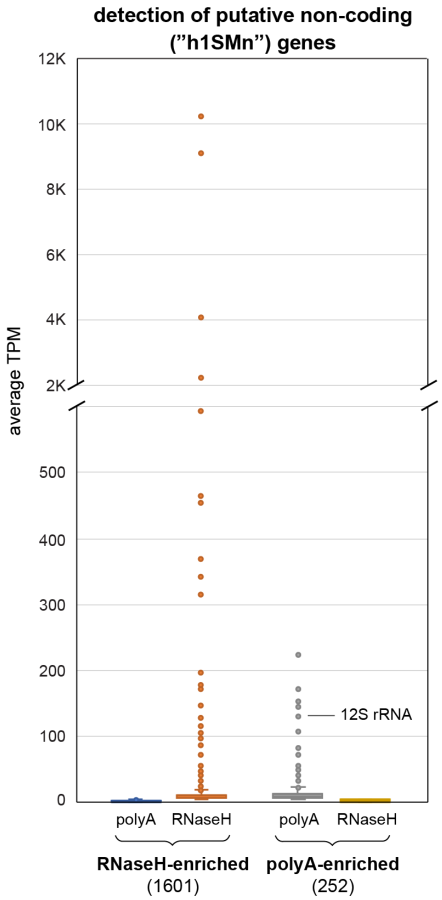


**Supplemental Figure S4.** Plot comparing detection of putative noncoding transcripts i.e., those starting with prefix h1SMn in the most recent set of S.med gene annotations (Ivankovic 2023) in polyA vs RNaseH-generated libraries. RNaseH-enriched = genes with TPM > 5 in RNaseH libraries but < 5 TPM in polyA libraries, polyA-enriched genes = genes with TPM > 5 in polyA libraries but < 5 TPM in RNaseH libraries.
